# Merlin Tumor Suppressor Function is Regulated by PIP_2_-Mediated Dimerization

**DOI:** 10.1101/2021.11.11.468247

**Authors:** Robert F. Hennigan, Craig S. Thomson, Kye Stachowski, Nicolas Nassar, Nancy Ratner

**Affiliations:** Division of Experimental Hematology and Cancer Biology, Cincinnati Children’s Hospital Medical Center, Department of Pediatrics, University of Cincinnati, Cincinnati, OH 45267-0713, USA

## Abstract

Neurofibromatosis Type 2 is an inherited disease characterized by Schwann cell tumors of cranial and peripheral nerves. The *NF2* gene encodes Merlin, a member of the ERM family consisting of an N-terminal FERM domain, a central α-helical region and a C-terminal domain that binds to the FERM domain. Changes in the intermolecular FERM-CTD interaction allow Merlin to transition between an open, FERM accessible conformation and a closed, FERM-inaccessible conformation, modulating Merlin activity. These conformational transitions are regulated by both phosphorylation and phosphoinositide binding. Merlin has been shown to dimerize, but the regulation and function Merlin dimerization is not clear. We used a nanobody based binding assay and found that Merlin dimerizes via a FERM-FERM interaction, orientated with each C-terminus in close proximity. Dimerization. Patient derived and structural mutants show that dimerization controls interactions with specific binding partners, including HIPPO pathway components, and correlates with tumor suppressor activity. We demonstrate that dimerization occurs after a PIP2 mediated transition from a closed to open conformation monomers that are then able dimerize. This process requires the first 18 amino acids of the FERM domain and is inhibited by phosphorylation at serine 518. The discovery that active, open conformation Merlin is a dimer represents a new paradigm for Merlin function with implications for the development of therapies designed to compensate for Merlin loss.

## Introduction

Neurofibromatosis Type 2 is an inherited disease characterized by schwannomas, benign Schwann cell tumors of cranial and peripheral nerves known as schwannomas. Schwannomas that arise in the general population also contain *NF2* mutations and represent 8% of all intracranial tumors ^1, 2^. Targeted deletion of the *Nf2* gene in mouse Schwann cells leads to schwannoma ^3, 4^ and *Nf2*-null cells have impaired contact inhibition of growth *in vitro* ^5, 6^. The *NF2* gene encodes Merlin, a 70-kDa member of the Ezrin-Radixin-Moesin (ERM) branch of the band 4.1 superfamily ^7, 8^. Merlin is predominately localized to the inner face of the plasma membrane ^9, 10^ where it associates with lipid rafts, which are membrane microdomains enriched with a variety of signaling molecules ^11^. Merlin is also a component of cell junctional complexes including adherens junctions and focal adhesions ^12, 13^. This is consistent with reports indicating that Merlin is a component of the HIPPO pathway, a growth suppressive signaling system which acts downstream of cell junctions and is responsible for contact inhibition of growth ^14–16^. Merlin loss also activates other oncogenic signaling networks, including Ras-ERK, Rac, src, β-catenin, and the mTOR protein kinase complex ^6, 17–26^. However, the mechanism by which these systems are activated is not fully defined.

Merlin has a conserved secondary structure consisting of an N-terminal FERM domain followed by a central α-helical (CH) region that folds over itself to form an anti-parallel coiled-coil that positions a C-terminal domain (CTD) for an intramolecular interaction with the FERM domain ^5, 27–29^. Merlin also has a unique N-terminal 20 amino acid sequence, absent in other ERM proteins, that is necessary for its tumor suppressor activity ^30^. The intramolecular interaction between the Merlin CTD and FERM domains mask a large portion of the FERM domain surface area, resulting in a closed, FERM-inaccessible conformation ^31^. Upon the release of the CTD from the FERM domain, Merlin transitions to an open conformation that renders the FERM domain accessible to critical interacting proteins ^32^. A Merlin mutant designed to stabilize the FERM-CTD interaction results in a constitutively closed conformation that has impaired tumor suppressor activity ^32^. Conversely, an open, FERM-accessible conformation mutant retains activity ^32^, demonstrating that Merlin’s tumor suppressor function is facilitated by the open conformation.

Merlin is phosphorylated at serine 518 by PAK2 ^33^ and PKA ^34^. This phosphorylation promotes a closed conformation with reduced Merlin tumor suppressor activity ^32, 35–37^. Merlin is also regulated by binding the lipid signaling molecule phosphatidylinositol 4,5 bisphosphate (PIP_2_) ^38^. This interaction promotes Merlin localization to the plasma membrane and is necessary for tumor-suppressive activity ^38^. PIP_2_ binding causes Merlin to assume an open, FERM accessible conformation ^39^ that is mediated by a structural change in the N-terminal portion of the central α-helical domain that forces the CTD and FERM domains apart ^40^. These reports show that Merlin activity is determined by conformational changes that are regulated by both phosphorylation and lipid based signaling systems. However, the specific mechanism by which open conformation Merlin interacts with binding partners to facilitate tumor suppression is not known.

The idea that Merlin forms dimers has long been established in the literature, initially by analogy to other ERM proteins and supported by the observation that its central α-helical domain may arrange itself into a coiled-coil structure ^41^. Experiments using yeast two-hybrid systems ^42, 43^ and GST pulldowns with isolated Merlin N-terminal FERM and C-terminal domains ^44–46^ suggested that Merlin exists as a dimer mediated by intermolecular FERM-CTD binding in an antiparallel orientation. However, other experiments using analytical ultracentrifugation explicitly ruled out dimerization ^39, 47^. Yet, a structure derived from co-crystalized Merlin FERM and CTD domains showed that Merlin dimerizes via interactions with between adjacent FERM from the dimerization partner ^48^. Given the contradictory nature of the literature, there is no clear consensus regarding the function of Merlin dimerization and its relationship to conformational regulation.

To address these questions, we used a nanobody based assay to measure Merlin binding in cotransfected cell lysates and between purified Merlin proteins ^13^. We rely on quantitative Merlin binding assays to assess dimerization. In addition, we use BRET as a corollary experimental system that allows us to define the relative orientation of one Merlin interacting with another. We find that Merlin dimerizes via a FERM-FERM interaction in a parallel orientation that requires the first 18 amino acids of the FERM domain. Patient derived and structural mutants show that the ability to dimerize correlates with tumor suppressor activity. Dimerization requires an open conformation, is inhibited by phosphorylation at serine 518 and is enhanced by PIP_2_ binding. These findings suggest novel mechanistic insights into Merlin function that integrate dimerization with conformation, phosphorylation and phosphatidylinositol signaling, providing novel mechanistic insights into Merlin function.

## Results

### Merlin Dimerizes via the FERM Domain

To investigate Merlin dimerization, we adapted a quantitative protein binding assay that utilizes a probe consisting of purified Merlin fused to the small, bright luciferase, NanoLuc, on its C-terminus. We used a high affinity anti-GFP nanobody bound to magnetic beads to isolate wild type Merlin with GFP fused to its C-terminus. The anti-GFP nanobody enabled the purification of Merlin-GFP fusion protein to homogeneity from transfected HEK 293T cells. Binding assays were performed by incubating the Merlin-NL probe with the purified Merlin-GFP bound to anti-GFP nanobody magnetic beads (Fig 1A) ^13^.

**Fig 1.**
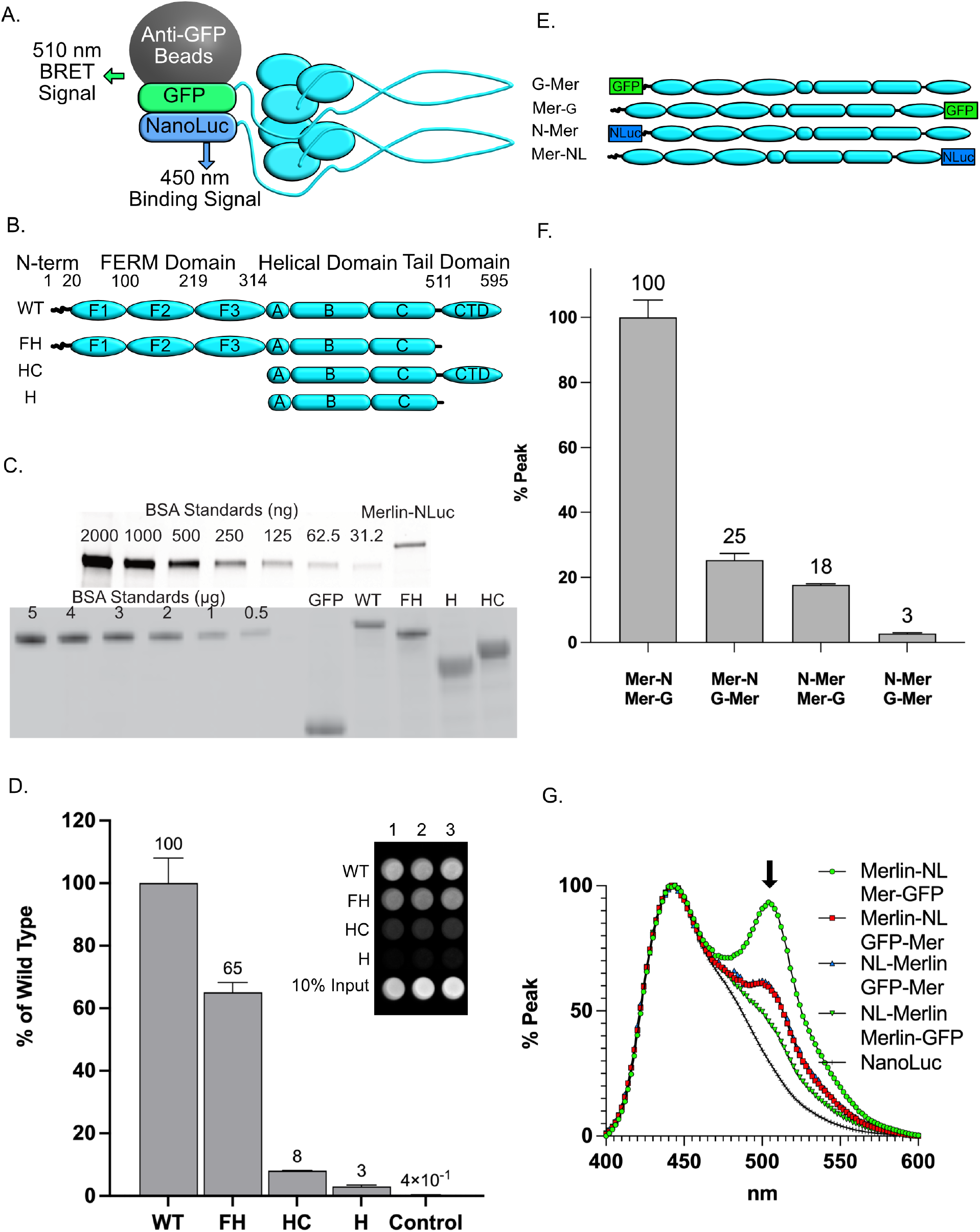
Merlin Dimerizes via the FERM Domain. **A).** A diagram depicting the dimerization assay using Merlin with either GFP or NanoLuc fused to its C-terminus. **B).** A schematic diagram depicting the full-length Merlin (aa 1-595, WT) and the C-terminal deletion mutant (FH, aa 1-511), the FERM domain deletion mutant (HC, aa 315-595) and the dual FERM and CTD deletion (H, aa 315-511). **C).** SDS-PAGE gels stained for total protein to access the quantity and purity of isolated probes for Merlin-NL (top) and GFP “bait” proteins (bottom) for GFP, Merlin-GFP, Merlin-FH-GFP, Merlin-HC-GFP and Merlin-H-GFP. **D).** Merlin dimerization data quantified on a plate reader and presented as the bound luciferase normalized to 10% of the unbound input NL probe. The data is a mean of triplicate binding reactions with standard deviation and expressed as a percentage of the wild-type Merlin. Inset: an image of the light emitted from Merlin-NanoLuc bound to Merlin and Merlin deletion mutant-GFP bound magnetic beads, performed in triplicate and dispensed into the wells of a 96-well plate (left). **E).** A schematic diagram depicting the N- and C-terminal GFP fused “bait” and the N- and C-terminal fused Merlin-NanoLuc “Probe” constructs used in the BRET assays. **F).** Triplicate Merlin dimerization assays for the N-and C-terminal Merlin constructs. The data is a ratio of bound to unbound NL probe, normalized to GFP fluorescence, (mean of triplicate binding reactions with standard deviation and expressed as a percentage of the peak value). **G).** Emission spectrum from 400 nm to 600 nm of Merlin dimerization assays normalized to the 450 nm peak. The BRET emission peak at 510 nm is indicated by the arrow.

To map the domains required for interaction we generated deletion mutants of the major Merlin domains, the FERM and CTD with C-terminal fusion of GFP and NanoLuc (Fig 1B). Purified Merlin-NL (Fig 1C) was incubated with Merlin-GFP bound beads (Fig 1C), washed then assayed for luciferase activity. Merlin dimerization was indicated by the amount of luciferase activity normalized to the amount of GFP-Merlin on the beads relative to control. We found that wild-type Merlin-NL bound to Merlin-GFP more than 100-fold greater than control GFP (Fig 1D). This experiment was conducted with the canonical Merlin isoform 1. Further binding experiments demonstrated that both Merlin isoforms 1 and isoform 2 dimerize with themselves and heterodimerize with each other (Supplemental Figure 1). Normalized binding data from Merlin deletion mutants revealed that the FERM domain is necessary and sufficient for dimerization (Fig 1D). The reduced binding to the FH mutant that lacks the CTD and the small amount of binding to the HC mutant containing the CTD is evidence of a relatively weak FERM-CTD interaction (Fig 1D). The absence of dimerization in the Merlin-H mutant shows that the central α-helical domain does not interact (Fig 1D). This experiment demonstrated that Merlin dimerizes primarily via a FERM-FERM interaction.

### Merlin Dimer Orientation

The luciferase substrate emission spectrum peaks 450 nm and is close enough to GFPs excitation peak at 495 nm to engage in bioluminescence resonance energy transfer (BRET), which occurs when NanoLuc and the GFP are less than 10 nm apart ^49^. Emission at 510 nm from Merlin-GFP:Merlin-NL complexes is evidence of BRET, indicating close proximity of the NanoLuc and the GFP proteins. We took advantage of this property to determine the orientation of the Merlin dimeric complex. HEK 293T cells were co-transfected with pairs of Merlin fused to GFP or NanoLuc at either at its N- or C-terminus, Merlin-GFP:Merlin-NL, GFP-Merlin:Merlin-NL, NL-Merlin:Merlin-GFP or GFP-Merlin:NL-Merlin (Fig 1E). The proximity of the N- and C-termini of each dimerization partner is measured by BRET, thus indicating the overall orientation of the dimer. Merlin complexes were immunoprecipitated and emission spectra from 400 nm to 600 nm were acquired. There was robust dimerization when GFP and NanoLuc are fused to the Merlin C-terminus (Merlin-GFP:Merlin-NL, Fig 1F). The BRET emission peak at 510 nm is apparent in Merlin-GFP:Merlin-NL, indicating close proximity of the GFP and NanoLuc proteins in this complex (Fig1G). When N-terminal and C-terminal fusion proteins were paired (NL-Merlin:Merlin-GFP or GFP-Merlin:Merlin-NL), significantly less dimerization was apparent (Fig 1F) and the 510 nm BRET peak was much reduced (Fig 1G). When both GFP and NanoLuc are N-terminal, dimerization was significantly reduced and the BRET signal was lost (GFP-Merlin:NL-Merlin, Fig 1E, D). Taken together, these data are consistent with the idea that Merlin forms a dimer. Dimer formation requires an uncovered N-terminus, and the complex is orientated such that each C-terminus is close to the other, consistent with a FERM-FERM interaction. Since the N-terminal fusion proteins show reduced dimerization, the following experiments were conducted with the C-terminal Merlin-GFP and Merlin-NL.

### Merlin Dimerization Correlates with the Open Conformation

The Merlin interaction assays and BRET assays strongly suggest that Merlin dimerizes. However, it remained possible that our data was the result of protein aggregation rather than dimerization. To exclude this possibility, we determined the size of purified Merlin-NL complexes using gel filtration chromatography (Fig 2A). Fractions from the column were assayed for NanoLuc activity to determine the Merlin-NL elution profile and the approximate size was calculated in reference to the elution profile of a set of standards of known Stokes radii (Supplemental Figure 2). Merlin-NL resolved into two peaks, a major peak with a Stokes radius of approximately 5.0 nm and a minor peak of 2.1 nm (Figure 2A). There was no evidence of NanoLuc activity above this peak at, ruling out Merlin-NL aggregation in these experiments. Total protein and western blots of fractions showed full length Merlin (Figure 2B), ruling out the possibility that the 2.1 nm peak is a result of proteolytic degradation. Merlin that is phosphorylated at S-518 was confined to the 5.0 nm peak (Figure 2B). To determine mobility of the Merlin dimer we co-expressed Merlin-GFP and Merlin-NL and then performed gel filtration chromatography on the cell lysates. Fractions from the column were assayed for NanoLuc activity then each fraction was immunoprecipitated with anti-GFP nanobody to determine the elution profile of the Merlin-GFP:Merlin-NL complex. GFP pulldowns from the fractions show that the Merlin-GFP:Merlin-NL complex elutes as a single peak at 5.0 nm (Figure 2C). There was minimal dimer detected above this peak, ruling out the possibility that dimerization is caused by non-specific aggregation. The total amount of NanoLuc activity pulldown from this peak was 2% of the total in the fraction. This small percentage of Merlin dimer is consistent with a Kd for dimerization being significantly higher than the nanomolar concentrations at which the chromatography was performed. The remaining 98% of the peak represents monomeric Merlin-NL.

**Fig 2.** Merlin Conformations. **A).** Fractionation of purified Merlin-NL (inset) on an SEC 650 gel filtration column, aliquots from 0.2 ml fractions were assayed for NanoLuc activity and plotted with the elution volume. The approximate Stokes radius of the peaks was calculated relative to the mobility of standards of known molecular weight and Stokes radius. (Supplemental Figure 1). The large peak at 12.4 ml has an approximate Stokes radius of 5.0 nm and the smaller peak at 15 ml has an approximate Stokes radius of 2.1 nm. **B).** SDS-PAGE with BSA standards and an aliquot of the purified Merlin-NL stained for total protein (top). Fractions composing peaks 1 and 2 were immunoblotted and probed with antibodies for either total Merlin or P-S518-Merlin (bottom). **C).** Gel fractionation of lysates from cells co-transfected with Merlin-NL and Merlin-GFP. NanoLuc activity from the fractions is shown as blue circles and depicted on the right-hand y-axis. Dimers represented by NanoLuc activity from GFP pulldowns of the fractions is shown as green squares and depicted on the left-hand y-axis. **D).** Gel fractionation of purified Merlin-NL at in hypotonic (50 mM NaCl), isotonic (150 mM NaCl) and hypertonic (500 mM NaCl) conditions. **E).** Images of dimerization assays dimerization assays performed at 50 mM, 75 mM, 150 mM, 300 mM and 600 mM NaCl. The GFP-NanoLuc fusion protein used to control for the effects of salt on luminescence. **F).** Quantitation of the dimerization data presented as a mean of triplicate binding reactions with standard deviation and expressed as a percentage of the peak value. **F).** Immunoblot assays of iHSC-1λ immortalized Schwann cell lysates fractionated by gel filtration and probed for either total Merlin (top) or P-S518-Merlin (bottom).

To determine if the 5.0 nm and 2.1 nm peaks represent different conformations of Merlin, we performed gel filtration experiments at different salt concentrations. Under isotonic conditions the ratio of the 5.0 nm to the 2.1 nm peaks was 2.6:1 (Fig. 2D). Under hypotonic conditions there was a shift from the 5.0 nm peak to the 2.1 nm peak with the ratio decreasing to 0.76:1 (Fig. 2D). Under hypertonic conditions the ratio further shifts to the 5.0 nm form, with a ratio of 3.6:1 (Fig 2D). These data suggest that the two peaks resolved by gel filtration represent two Merlin conformations. Dimerization assays performed at different salt concentrations showed that Merlin-Merlin binding was maximal at low salt (50 mM) and lost at high salt (300 mM and 600 mM, Fig 2E, F), suggesting that the ability to dimerize correlates with the 2.1 nm peak. To confirm that the conformers are not an artifact of overexpression in HEK cells, we performed gel filtration experiments on cell lysates from immortalized human Schwann cells and assayed fractions for endogenous Merlin by western blot (Fig.2G). The majority of endogenous Merlin elutes with peak 1 at 13 ml (5.0 nm) with a small amount in a fractions that elute with the 2.1 nm peak at 15 ml (2.1 nm). As with the recombinant Merlin, phospho-518 Merlin was restricted to the 5.0 nm peak.

### Merlin Mutant Dimerization

We evaluated the ability of a panel of Merlin mutants to dimerize (Fig 3A). The patient derived mutants Δ39-121, L360P and L535P all showed significantly reduced dimerization relative to wild type (Fig 3B). However, the patient derived L64P mutant showed slightly enhanced dimerization (1.28-fold, Fig 3B); suggesting that this mutant interferes with aspects of Merlin function that are independent of dimerization. Of structural mutants, a lipid binding deficient mutant 6N had impaired dimerization (Fig 3B). A deletion of the Merlin unique N-terminal 18 amino acids (ΔN18) showed significantly impaired dimerization, 33% relative to wild type (Fig 3C). However, GFP fused to the first 20 N-terminal amino acids (N20) failed to dimerize (Fig 3C) indicating that Merlin’s unique 18 amino acid N-terminus is necessary but not sufficient for full dimerization. A non-phosphorylatable mutant at serine 518, S518A, had significantly enhanced dimerization whereas its phosphomimetic counterpart, S518D, had reduced dimerization (Fig 3C). This suggests that dimerization can be regulated by phosphorylation. A mutant designed to force Merlin into a constitutively closed conformation, Merlin-AR ^32^, failed to significantly dimerize relative to wild type. The constitutively open conformation mutant, Merlin-ΔEL ^32^, had significantly increased dimerization (Fig 3C).

**Fig 3.**
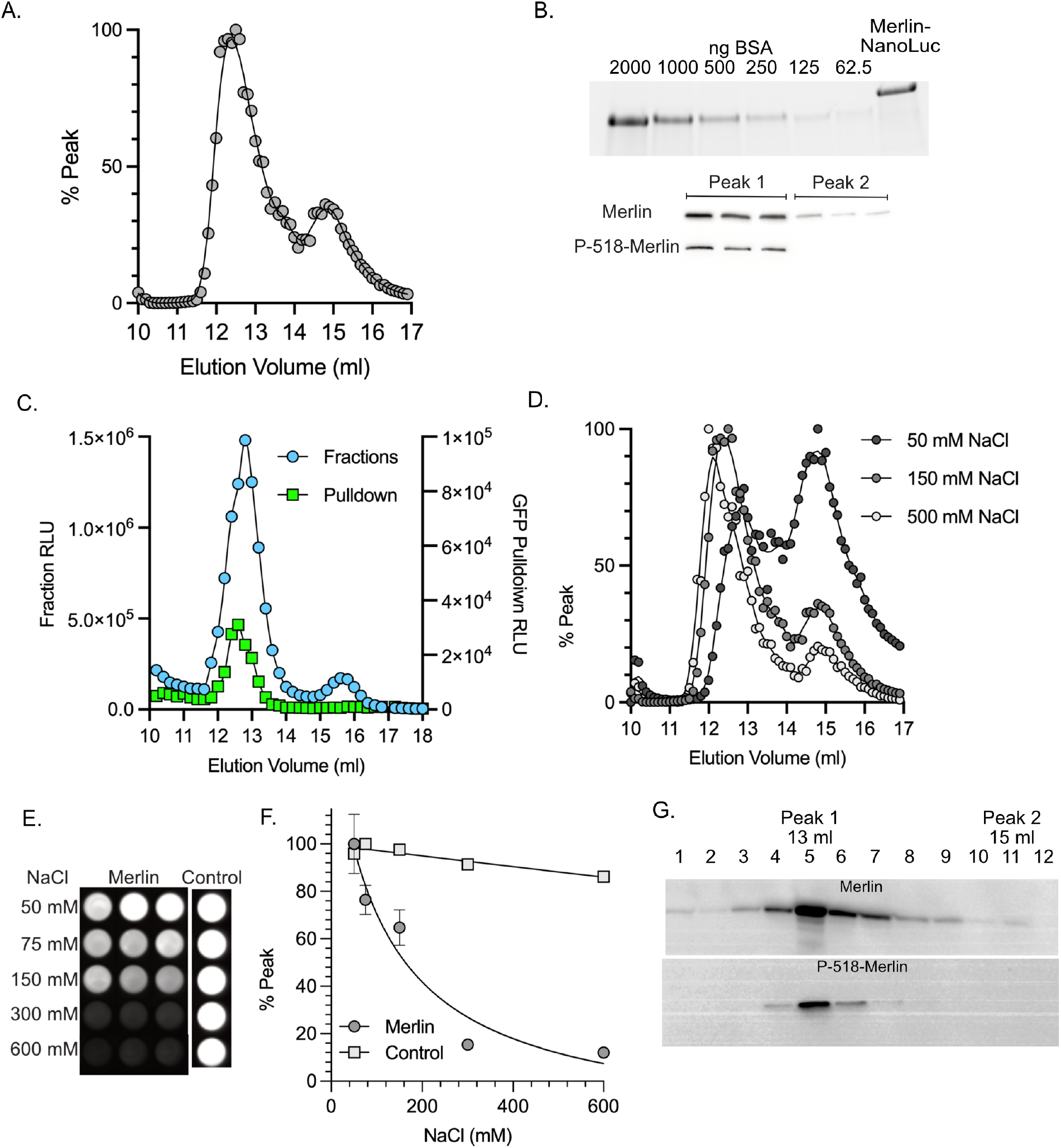
Merlin Mutant Dimerization. **A.** A schematic diagram of Merlin mutants. Patient derived mutations: L64P, L360P and L535P, indicated by asterisks. ΔN18: deletion of the N-terminal 18 amino acids. N20: the first 20 amino acids fused to GFP. S518A/S518D: non-phosphorylatable and phosphomimetic mutants at S518. AR: closed conformation mutant. ΔEL: open conformation mutant. **B).** Pulldown dimerization assays of patient derived Merlin mutants, L64P, Δ39-121, L360P, L535P and the lipid binding deficient mutant 6N. **C).** Pulldown dimerization assays of Merlin N-terminal mutant, ΔN18, the unique N-terminal 20 amino acids fused directly to GFP, N20, the phosphorylation mutants S518A and S518D and the conformation mutants AR and DEL. GFP alone serves as the negative control. The data is a ratio of bound to unbound NL probe, normalized to GFP fluorescence. (mean of triplicate binding reactions with standard deviation and expressed as a percentage of the wild type Merlin. **D).** Gel fractionation of lysates from cells transfected with plasmids expressing either wild type Merlin-NL (blue circles) Merlin-S518D-NL (red squares) or Merlin-S518A-NL (green triangles). **E).** Gel fractionation of lysates from cell transfected with plasmids expressing either wild type Merlin-NL (blue circles) Merlin-AR-NL (red squares) or Merlin-ΔEL-NL (green triangles).

We tested if dimerization of these Merlin mutants correlates with changes in conformers, as assessed by gel filtration. These experiments showed that the S518D mutant had a reduced 2.1 nm peak relative to both wild type and S518A (Fig 3D). This is a further correlation between the small peak and the ability to dimerize and is consistent with the western blots in Figure 1 showing that phospho-518 Merlin is restricted to the large peak. Gel filtration experiments showed that the closed conformation Merlin-AR mutant failed to form the small 2.1 nm peak relative to both wild type and the open conformation Merlin-ΔEL mutant (Fig 3E). These data suggest that the 2.1 nm peak represents an “open” conformation monomer and that the ability to form the open conformer correlates with the ability to dimerize. Together these data show that Merlin dimers are hypo-phosphorylated and in the open conformation. BRET studies confirmed these results, indicating a tighter interaction between S518A dimers than S518D dimers and a similar tight interaction is indicated by the greater BRET signal in Merlin-ΔEL relative to Merlin-AR (Supplemental Fig 3).

### Interaction of Dimerization Mutants with Merlin Targets

We hypothesized that Merlin dimerization might affect its interaction with known target proteins. To test this, we performed co-transfection pulldown experiments with plasmids expressing wild type or mutant Merlin-NL and GFP fused to each of four known Merlin binding proteins: Angiomotin, ASPP2, Lats1 and YAP1 ^13, 50–52^. All four proteins showed significantly reduced binding to the phosphomimetic S518D mutant relative to S518A (Fig 4A-D). Angiomotin bound to the constitutively closed Merlin-AR mutant but showed impaired binding to the open conformation mutant Merlin-ΔEL (Fig 4A). In contrast, ASPP2, Lats1 and YAP1, all showed either equivalent or significantly increased interaction to Merlin-ΔEL mutant relative to Merlin-AR (Fig 4B-D). These experiments suggest that dimerization reduces the interaction of Merlin with Angiomotin and increases interaction with the key target proteins ASPP2, Lats1 and YAP1

**Fig 4.**
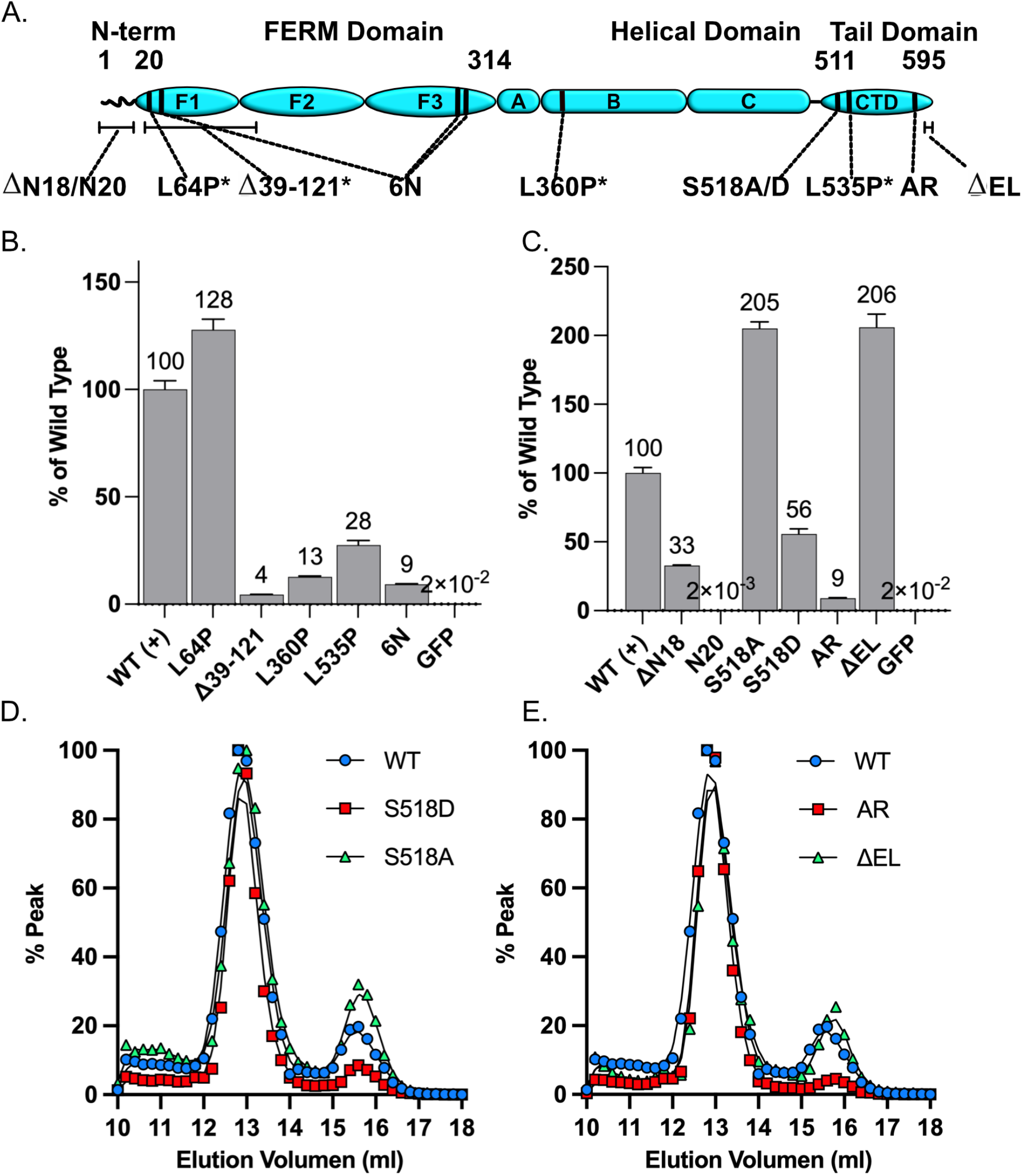
Interaction of Dimerization Mutants with Merlin Targets. Normalized luciferase activity of GFP pulldowns from HEK 293T cells transfected with plasmids expressing wild type and mutant Merlin-NL and GFP fusions for **A)**. Angiomotin, **B)**. ASPP2, **C).** YAP1, **D).** Lats1. The data is a ratio of bound to unbound NL probe, normalized to GFP fluorescence shown as a mean of triplicate binding reactions with standard deviation and expressed as a percentage of the wild-type Merlin. Asterisks indicate significance by a two-way ANOVA test).

### Effect of PIP_2_ and Phosphorylation on Dimerization

Crystallographic experiments showed that upon binding to the signaling phosphoinositide, PIP_2_, Merlin assumes an open conformation ^39, 40^. We confirmed this using gel filtration experiments. In the presence of 200 μM the soluble PIP_2_ analog PIP_2_-diC8, Merlin shifted to the 2.1 nm open conformer (Fig 5A). Since the open conformation correlates with dimerization, we tested if PIP_2_ binding also caused enhanced dimerization. We performed binding assays with purified Merlin-GFP and Merlin-NL in increasing concentrations of. This experiment showed increased dimerization from 6.25 to 400 μM PIP_2_-diC8, with a plateau above 200 μM (Fig 5B).

**Fig 5.**
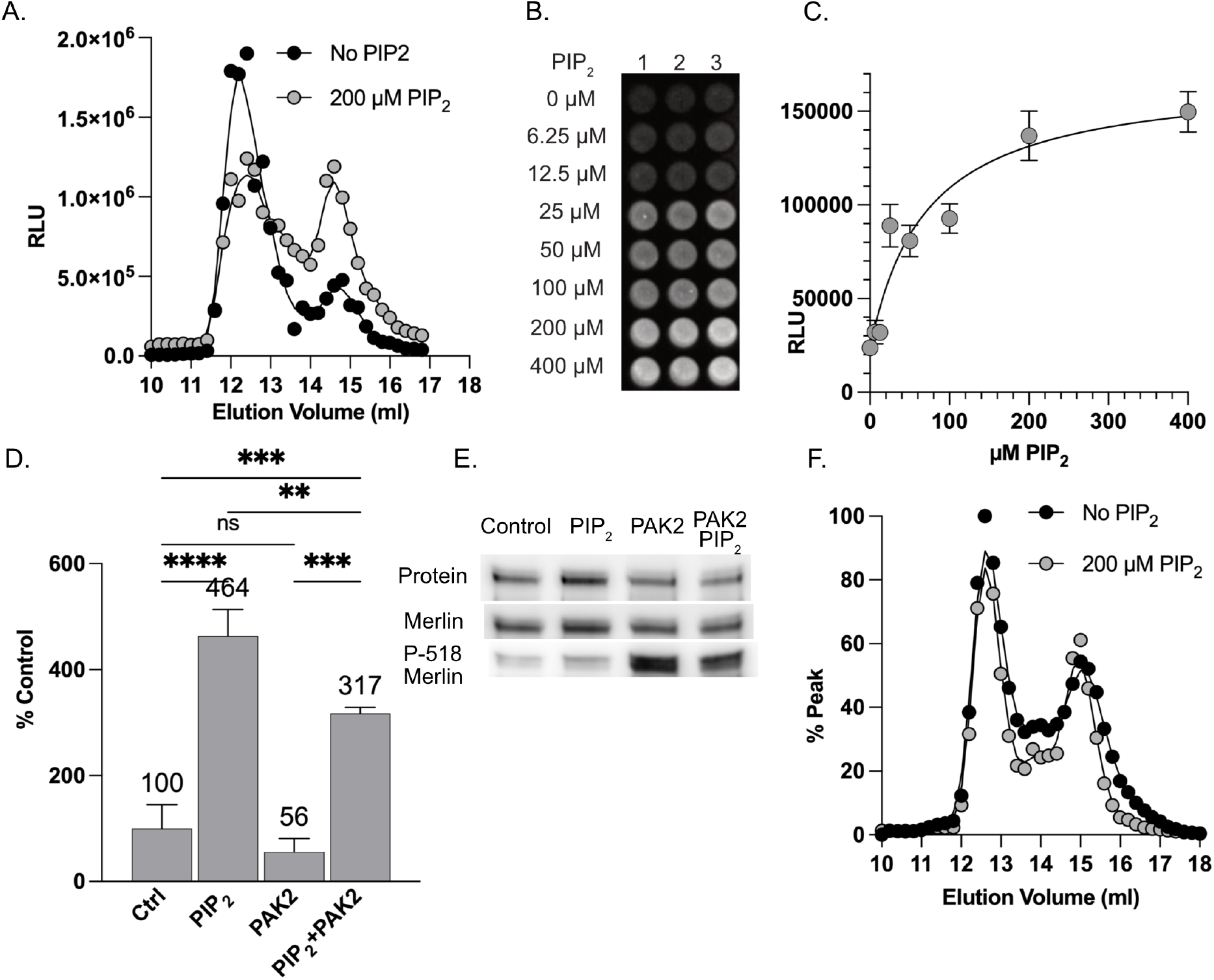
Effect of PIP_2_ and Phosphorylation on Dimerization. **A).** Gel fractionation of purified Merlin-NL in the absence (black circles) or presence (grey circles) of 200 μM PIP_2_-DiC8. **B).** An image of the dimerization assay using 100 nM purified Merlin-GFP and Merlin-NL incubated with increasing concentrations of PIP_2_-DiC8. **C).** Binding curve of Merlin dimerization data expressed in relative luciferase units (RLU) and fitted to a binding curve. **D).** Dimerization assays using each of purified Merlin-GFP and Merlin-NL in with and without 200 μM PIP_2_-DiC8 and/or *in vitro* phosphorylation by recombinant PAK2. The data is presented as the bound luciferase normalized to 10% of the unbound input NL probe. The data is a mean of triplicate binding reactions with standard deviation and expressed as a percentage of the untreated control. **E).** Western blots of proteins recovered from the dimerization assay showing total protein (top), probed with antibodies to Merlin (middle) or P-518 Merlin (bottom). **F).** Gel fractionation of purified Merlin-DN18 NanoLuc in the absence (black circles) or presence (grey circles) of 200 μM PIP_2_-DiC8.

**Fig 6.**
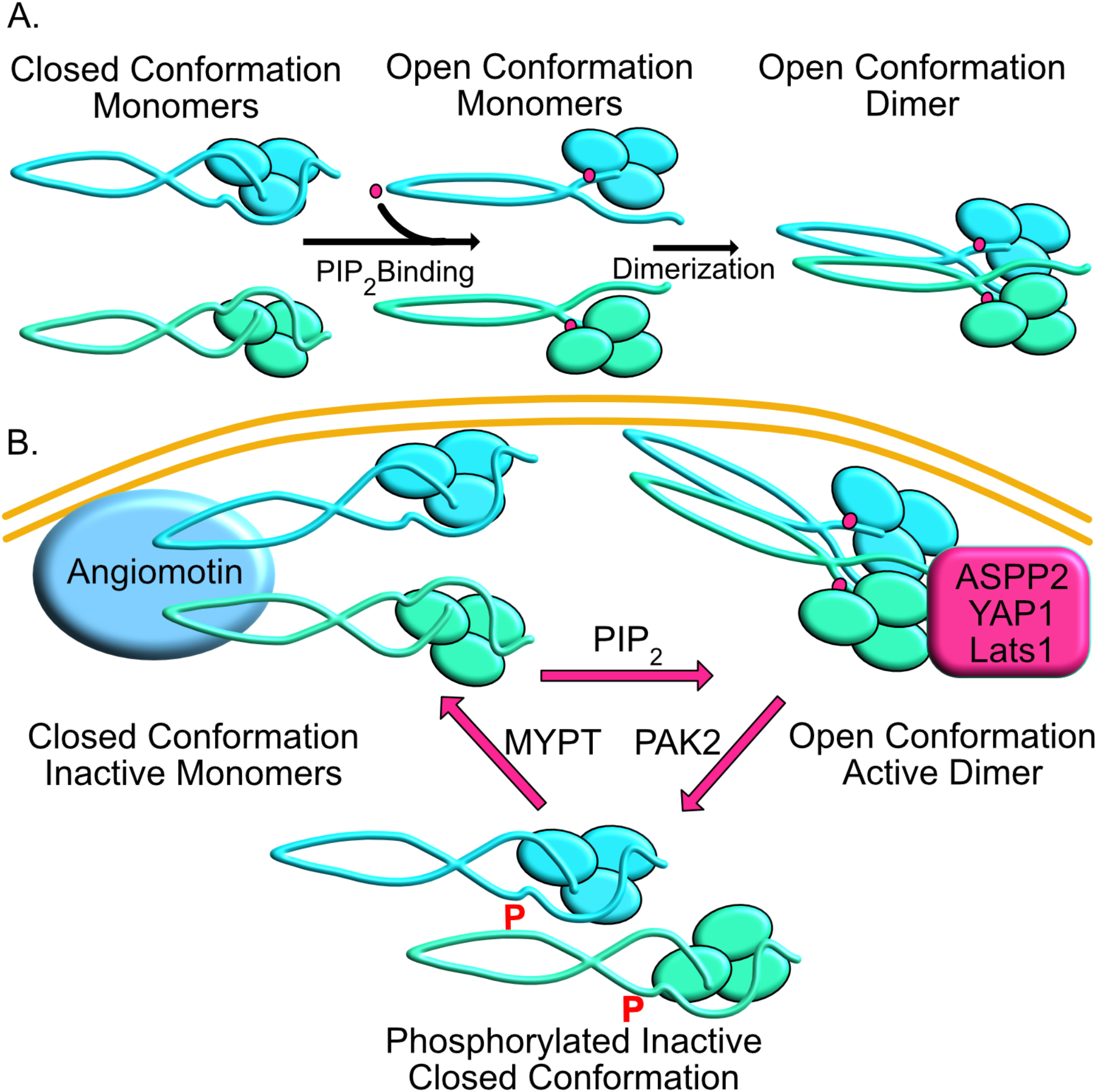
Merlin Activation Cycle. **A).** Model for PIP_2_ mediated Merlin dimerization with “closed” monomers transitioning to “open” monomers upon PIP_2_ binding the proceeding to dimerization. **B).** Model for Merlin activation cycle. Inactive closed conformation associated with Angiomotin. PIP_2_ binding leading to open conformation, dimerization and binding to ASPP2/Lats1/YAP1. Active complexes are phosphorylated leading to dissolution of the complex followed by dephosphorylation to regenerate the Merlin-Angiomotin complex.

Since the phosphomimetic S518D mutant had impaired dimerization relative to non-phosphorylatable S518A, we tested if *in vitro* phosphorylation inhibited dimerization in the presence and absence of PIP_2_-diC8. Merlin dimerization was increased in the presence of PIP_2_-diC8 by 4.64-fold and phosphorylation by PAK2 decreased the PIP_2_ effect significantly (Fig 5C). Western blots confirmed increased levels of phospho-S518 in the PAK2 treated samples (Fig 5D). These results suggest a regulatory cycle in which Merlin is activated by PIP_2_, causing it to assume an open conformation that allows dimerization, leading to interactions with target proteins such as ASPP2, Lats1 and YAP1. This “active” tumor suppressing complex is then dismantled upon phosphorylation at S518 by PAK2.

Since we found that the ΔN18 N-terminal deletion mutant is deficient in dimerization and, like the 6N mutant, this region has been shown to localize Merlin, to bind lipids and to localize to the plasma membrane ^30^, we ran gel filtration experiments to test for PIP_2_-mediated conformational change. Although the ΔN18 mutant can assume the open conformation it failed to undergo the shift to the open conformation like wild type does (Fig. 5F). This suggests that a primary function for Merlin’s unique N-terminus is to facilitate the PIP_2_ mediated shift to the open conformation.

## Discussion

We demonstrate that Merlin dimerizes via a FERM-FERM interaction, in a parallel orientation and that dimerization requires an uncovered N-terminus and the first 18 amino acids of the FERM domain. Our data is inconsistent with the prevailing model of ERM dimerization that posits an intermolecular FERM-CTD interaction between adjacent Merlin molecules ^43, 53^. However, there is precedent in the literature for FERM domain mediated Merlin dimerization ^48^. A structure derived from co-crystalized Merlin FERM and CTD domains showed a Merlin dimer with extensive interactions between FERM domains, mediated by unfurling the F2 lobe of the FERM domain such that it interacts with an adjacent FERM from the dimerization partner ^48^. This structure is consistent with the FERM mediated dimerization that we describe.

Our gel filtration experiments revealed that Merlin exists in two populations that represent predominately monomeric closed and monomeric open conformations. Dimeric Merlin is detectable as a small minority of this population, roughly 2%, that co-elutes with the closed monomeric conformation. The small percentage of dimers in this population is a function of the low, nanomolar concentrations at which the binding assays are conducted. The low binding percentage at these concentrations would suggest a much higher Kd for the Merlin dimerization reaction, likely to be in the micromolar range. The behavior of the conformation mutants, Merlin-AR and Merlin-ΔEL, show that the open conformation is necessary for dimerization. The inability of Merlin-AR to dimerize is telling. Merlin-AR is an inactivating mutant that has amino acid substitutions in the extreme C-terminus, A585W and R588K, that stabilize the intramolecular FERM-CTD interaction ^32^ thus fixing Merlin into the “closed’ conformation that we find is unable to dimerize. In contrast Merlin-ΔEL has a deletion of the last two C-terminal amino acids, E594 and L595, that weakens the intramolecular FERM-CTD interaction leading to a more “open” conformation ^32^. This mutant retains tumor suppressor activity ^32^ and has enhanced dimerization relative to wild type, again linking dimerization with the open conformation and tumor suppressor function. Also, Merlin-ΔEL shows enhanced binding to three of four Merlin binding proteins, ASPP2, YAP1 and Lats1. In contrast, binding of S518D with Angiomotin, ASPP2, YAP1 and Lats1 is reduced relative to both S518A and wild type. This enhanced binding suggests that the open conformation Merlin dimers represent an activated state with increased affinity for critical target proteins.

We show that PIP_2_ binding to Merlin causes a shift to the monomeric open conformation and that Merlin’s unique N-terminus is necessary to shift to the open conformation in the presence of PIP_2_. Therefore, this small domain is a critical point of regulation by phospholipid second messengers. This region contains basic amino acids that are necessary for Merlin mediated growth suppression and localizes Merlin to the inner face of the plasma membrane ^30^ where it is a target of regulatory kinases that control Merlin stability ^37, 54^. These data are supported in the literature by a small angle neutron scattering study demonstrating that wild type Merlin exists in a closed conformation in the absence of PIP_2_ but in an open conformation in the presence of PIP_2_ ^39^. This is consistent with the crystal structure of PIP_2_ bound Merlin that shows a dramatic conformational change in the central α-helical domain that forces the CTD away from the FERM domain resulting in an open, FERM-accessible conformation ^40^. This conformational change results in increased affinity for the Merlin binding protein Lats1 ^40^. This is consistent with our finding that open conformation, dimerization mutant Merlin-ΔEL has increased binding for ASPP2, YAP1 and Lats1. Phosphorylation of Merlin at serine 518 correlates with the closed conformation and inhibits dimerization. Unlike wild type, the S518D mutants fail to adopt an open conformation upon PIP_2_ binding ^39^. Consistent with all these findings, we show that binding of S518D with Angiomotin, ASPP2, YAP1 and Lats1 is reduced relative to both S518A and wild type.

Together these results indicate that PIP_2_ bound, dimeric Merlin complexes are responsible for binding to critical target proteins that mediate tumor suppressor function. This suggests a regulatory loop in which hypo-phosphorylated Merlin is activated by PIP_2_, causing a shift to the monomeric open conformation followed by dimerization and interaction with binding partners. This active dimeric complex is then phosphorylated on serine 518 by PAK2 leading to a closed, inactive monomeric conformation. Merlin is a mechanosensory signaling molecule that is activated in response to specific biochemical signals that originate from cell junctions in response to extracellular mechanical cues ^55, 56^. For instance, PIP_2_ is generated at sites of N-cadherin–mediated intercellular adhesion as cells become confluent ^57, 58^, this could provide a locally focused stimulus for Merlin to first undergo conformational change and then dimerize. Activated Merlin may then bind to specific target proteins such as Angiomotin, Lats1 and YAP1 through which it regulates the HIPPO pathway ^15, 59–62^. PIP_2_ mediated dimerization thus provides a mechanism that allows local activation of Merlin and binding to target proteins at a specific subcellular location.

Ultimately, understanding the biochemical mechanism of Merlin function will give insight into the consequences Merlin loss and is therefore critical to the development of novel strategies to treat NF2. The model we propose for Merlin function may impact potential therapies. For example, one strategy to treat NF2 proposes to use gene therapy to replace lost Merlin function in schwannoma cells. The ability of Merlin to dimerize raises the possibility that some mutant Merlin proteins may act as dominant negative or dominantly activating proteins, impairing the activity of ectopic wild type Merlin, potentially lessening effect of added wild type alleles. Secondly, it is clear from proximity biotinylation experiments ^13^ that Merlin may bind many other critical target proteins beyond the well described HIPPO pathway members Lats1, YAP1 and Angiomotin. Each novel binding partner represents a possible target for the development of drugs to treat Neurofibromatosis Type 2. It will be important to identify those novel binding partners that mediate Merlin’s tumor suppressor activity. It is now necessary identify target proteins that bind to Merlin in the active, PIP_2_-bound dimeric state, thus identifying different pathways that may lead to the development of new therapies to treat NF2.

## Materials and Methods

### Plasmids

pEGFP-C3-hYAP1 and pEGFP C3-Lats1 (Addgene plasmid # 19053) were gifts from Marius Sudol (Addgene plasmid # 17843). HA-AMOT p130 was a gift from Kunliang Guan (Addgene plasmid # 32821) pcDNA3.1-ccdB-Nanoluc was a gift from Mikko Taipale (Addgene plasmid # 87067). GFP-ASPP2 was a gift from Charles D. Lopez.

### Cloning

Recombinant expression plasmids were constructed by PCR amplification of both vector and insert sequences using Q5^®^ High-Fidelity DNA Polymerase (New England Biolabs, Ipswich, MA) followed by a Gibson assembly reaction using NEBuilder HiFi DNA Assembly Cloning Kit, then transformed into NEB-a Competent Cells (New England Biolabs, Ipswich, MA). A full listing of the primer sequences used to generate all the plasmids used in this study is provided in the Supplemental file. To facilitate assembly, primers were designed with 10-15 bp extensions that are homologous to regions flanking the in-frame insertion sites in the plasmids. These sites are indicated in the maps of the vectors and shown in the supplemental files. All primers were purchased from Integrated DNA Technologies, Inc. (Coralville, IA).

The expression vectors used in the study were initially derived from pEGFP-N1 and pEGFP-C1 (Takara, La Jolla, CA). A double stranded oligomer encoding StrepTag II affinity tag was cloned, in-frame, N-terminal to EGFP in pEGFP-C1 and C-terminal to EGFP in pEGFP-N1 to generate pST-EGFP and pEGFP-st, respectively. NanoLuc insert was amplified from pcDNA3.1-ccdB-Nanoluc, inserted in frame with the StrepTag II in place of EGFP in pST-EGFP and pEGFP-st to generate pST-NanoLuc and pNanoLucST. Maps and insertion sites of these plasmids are shown in the supplemental file. Full-length Merlin isoform 1 was PCR amplified form pC-Mer-Vst ^63^ to generate pMer-GFPst pMer-NLst, pST-GFP-Mer and pST-NL-Mer. Similarly, Angiomotin was PCR amplified from pHA-AMOT p130 and assembled into pST-EGFP to generate pST-GFP-AMOT. Merlin mutants were introduced into pMer-GFPst and pMer-NLst Via PCR amplification and Gibson assembly using the primers listed in the supplemental file.

### Cell Culture

HEK-293T were used for co-transfection experiments and to purify Merlin-NL and Merlin-GFP proteins because they are readily transfectable, grow to high density and maintain SV40 ori containing plasmids in an episomal state. These features generate high yields of transfected recombinant proteins while utilizing endogenous mammalian chaperones and post-translational modification systems. The cells were purchased from ATCC (CRL-2316) grown in DMEM supplemented with 10% FBS and Pen/Strep at 37°C 5% CO_2_.

Immortalized human Schwann cell line, iHSC-1λ were a gift from Dr. Margaret Wallace (University of Florida College of Medicine, Gainesville, FL, USA)^64^ also grown in DMEM supplemented with 10% FBS and Pen/Strep at 37°C 5% CO_2_.

### Protein Purification

To purify Merlin-NL probes, 5 x 10^6^ HEK 293T cells were plated into each of two 15 cm dishes then transfected with 50 μg pMerlin-NLst plasmid per plate using PEI at a 3:1 PEI to DNA ratio. Cells were harvested in 0.5 ml TBS pH 8.0, 2 mM MgCl_2_, 1% CHAPS plus HALT protease inhibitor mix (Thermo Fisher Scientific, Waltham, MA) and 2.5 U Benzonase (Sigma Aldrich, St. Louis, MO), the lysates cleared by centrifugation at 20 kxG, 4°C for 10 min and supernatants were recovered. Washed magnetic Streptactin beads (100 μl, MagStrep type 3 XT, IBA Life Sciences, Gottingen, Germany) were added and incubated overnight at 4**°**C with agitation. Beads were recovered by magnet, washed 3x each with high salt buffer (20 mM Tris-Cl pH 8.0, 0.5 M NaCl, 1% Triton X-100, 0.05% Tween 20 + 0.5 mM EDTA), low salt buffer (20 mM Tris-Cl pH 8.0, 50 mM NaCl, 10% Glycerol, 0.5% NP-40, 0.05% Tween 20 + 0.5 mM EDTA) and an isotonic buffer (20 mM Tris-Cl pH 8.0,150 mM NaCl, 0.05% Tween 20 + 0.5 mM EDTA) then eluted in 200 μl elution buffer (100 mM Tris-Cl pH 8.0,150 mM NaCl + 50 mM D-biotin). Protein concentration and purity of an aliquot was evaluated by 4-20% SDS-PAGE against a standard curve of BSA stained with Instant Bands fluorescent total protein stain and visualized on an Azure C-600 imaging system (Dublin, CA).

For small scale affinity purification, 20 μg Merlin-GFP expressing plasmid was transfected into 2 x 10^6^ 293T cells in a 10 cm dish using PEI, lysed as described above then affinity purified with 40 μl washed GFP-Trap_MA beads (Bulldog Bio, Portsmouth), NH) overnight at 4°C with agitation. The beads were washed as described above and resuspended in TBST.

### Direct Binding Assays

The GFP bound beads were resuspended in a 30-μl blocking buffer of 0.5 mg/ml BSA in TBST and incubated for 1 hour at room temp., The beads were recovered magnetically then resuspended blocking buffer plus 30 to 50 nM Merlin-NL protein and incubated at room temperature for 1 hour. Beads were then recovered, washed four times with TBST the resuspended in 25 μl TBST. Luciferase activity was measured using 25 μl NanoGlo Luciferase Substrate Buffer (Promega, Madison, WI). NanoLuc activity, GFP fluorescence and BRET spectra were measured on a FlexStation 3 microplate reader (Molecular Devices, San Jose, CA). Binding assays were performed in triplicate. Binding is calculated as bound luciferase activity normalized to a 10% aliquot of the unbound input (*Bound* ÷ 10% *Input*). These values were expressed relative to WT. For visual reference, after counting on the plate reader, the luminescence in the microplates was imaged with an Azure C-600 imaging system (Dublin, CA).

### Co-Immunoprecipitation

HEK 293T cells were co-transfected with GFP-target and Merlin-NL plasmids, lysed in 0.5 ml of TBS, 2 mM MgCl_2_, 0.5% NP-40, protease inhibitor mix. Lysates were diluted 1:2.5 with TBST then NL activity was measured. GFP fusion proteins were immunoprecipitated using GFP-Trap_MA (ChromoTek, Hauppauge, NY), recovered using a magnetic stand, and washed four times with TBS. Luciferase activity was measured and imaged as described above. To control for GFP loading, fluorescence at 485 nm excitation and 510 emission was measured on the FlexStation 3. The binding is calculated as the ratio of bound luciferase activity to unbound luciferase activity (*Bound* ÷ *Unbound*)/*GFP FLuorescence*). Experiments were performed in triplicate and presented a s % of WT. Statistical significance was assessed by two-way ANOVA using Prism.

### Gel Filtration

Gel filtration chromatography was performed on a BioRad NGC Quest 10 Plus FPLC system, equipped with a BioFrac fraction collector, controlled by ChromLab software and using a 10 x 300 mm Enrich SEC 650 gel filtration column with a 24 ml bed volume. For each run the column was equilibrated with 2 volumes of buffer at 1 ml/min, 0.25 ml sample was then injected automatically, and 10 ml buffer was run through the column before 40, 0.2 ml fractions were collected for another 8 ml. Typical running buffer was filtered and degassed PBS. For the experiments using different salt conditions, two pumps were run with Buffer A; 20 mM Tris-Cl pH 7.4, 1 mM EDTA and Buffer B; 20 mM TrisCl pH 7.4, 500 mM NaCl, 1 mM EDTA. Hypotonic buffer at 50 mM NaCl was achieved by mixing 90% Buffer A and 10% Buffer B. Isotonic buffer at 150 mM NaCl was achieved by mixing 70% Buffer A and 30% Buffer B. Hypertonic buffer at 500 mM NaCl was achieved by mixing 0% Buffer A and 100% Buffer B. Samples were passed through a 0.22 μ spin filter (Ultrafree-MD. Millipore) before injection. Samples consisted of 5-20 μg purified Merlin-NanoLuc protein in 0.25 ml column buffer. Alternatively, cleared CHAPS lysates of cells expressing Merlin-NanoLuc constructs were desalted into PBS with PD MiniTrap G-25 desalting columns (Cytiva) and concentrated to 0.25 ml by a PES spin concentrator with a 10K MWCO (Pierce). NanoLuc assays were performed with a 10 μl aliquot of fractions transferred to opaque round bottom 96 well plates and luciferase activity was measured using 10 μl NanoGlo Luciferase Asssay reagent (Promega, Madison, WI) and luminescence was measured on a FlexStation 3 microplate reader (Molecular Devices, San Jose, CA). Graphs presenting luciferase activity (RLU) in each fraction or luciferase activity normalized to the activity in the peak fractions (%) relative to the elution volume in ml were prepared using Graphpad Prism 9 software.

### PIP_2_ and PAK

Active PAK2 (Sigma-Aldrich) was incubated with purified Merlin-GFP bound beads and Merlin-NL in 25 mM Tris (pH 7.4), 2 mM DTT, 10 mM MgCl_2_, 200 μM ATP, 150 mM NaCl_2_, +/- 200 μM PiP_2_-diC8 (Eschelon Biosciences), +/- 0.278 U/μl PAK2 for 1 hour at 30°C. Luciferase activity and BRET was measured as described above. Antibodies used for immunoblots were Phospho-Merlin (Ser518) (D5A4I), a rabbit mAb raised against synthetic peptide corresponding to residues surrounding Ser518 of human Merlin protein from Cell Signaling Technology Cat # 13281 and a mouse monoclonal antibody, 4B5, raised against recombinant Merlin ^47^. Both were used at a 1:1000 dilution.

## Acknowledgments

The authors would like to thank Dr. Shyra Tedesco and Dr. Brad Ozanne for stimulating discussion and critically reading the manuscript.

