## Supplemental Figure 1 for "Merlin Tumor Suppressor Function is Regulated by PIP_2_-Mediated Dimerization"

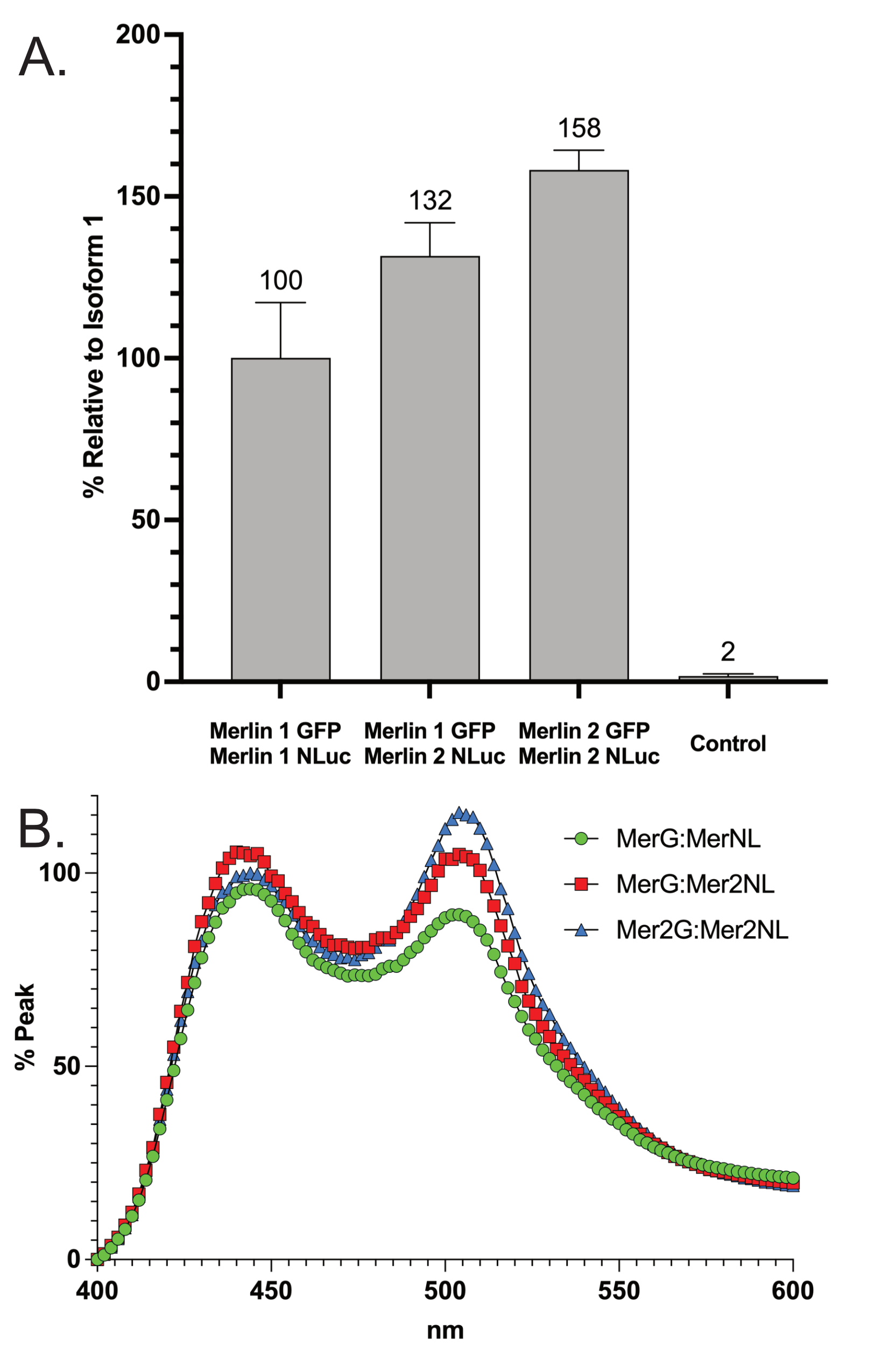

### Supplemental Figure 1.

A). Merlin isoform dimerization reactions with Merlin isoform 1-NL: Merlin isoform 1-GFP, Merlin isoform 2-NL: Merlin isoform 1-GFP and Merlin isoform 2-NL: Merlin isoform 2-GFP. The data is a mean of triplicate binding reactions with standard deviation and expressed as a percentage of the Merlin isoform 1.

B). Emission spectrum from 400 nm to 600 nm of Merlin isoform dimerization assays normalized to the 450 nm peak.
