## Supplemental Figure 2 for "Merlin Tumor Suppressor Function is Regulated by PIP_2_-Mediated Dimerization"

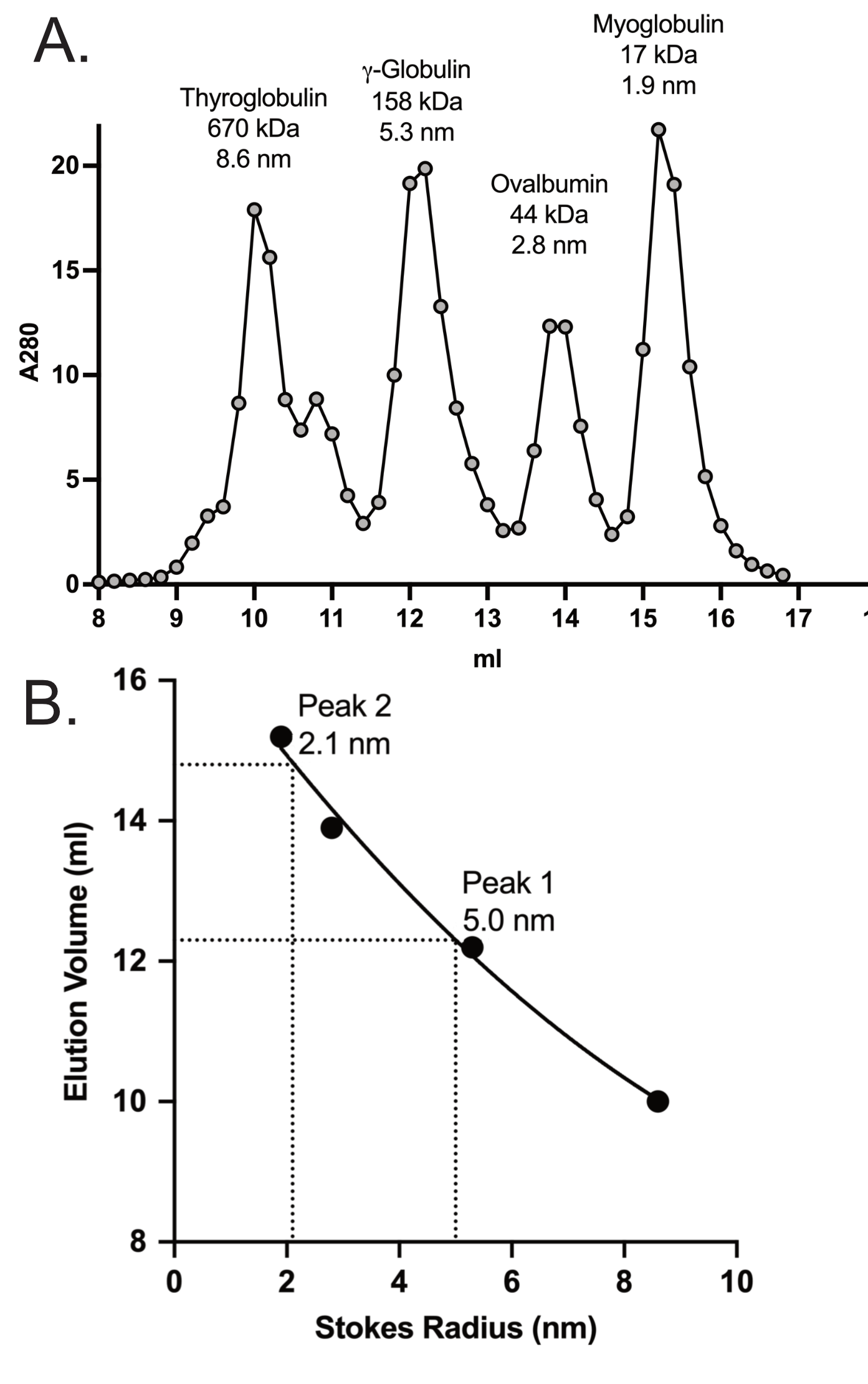

A). The A280 elution profile of 300 µg gel filtration standards run in PBS. The standards are thyroglobulin at 670 kDa with an 8.6 nm Stokes radius, γ-globulin at 158 kDa with a 5.3 nm Stokes radius, ovalbumin at 44 kDa with a 2.8 nm Stokes radius and myoglobulin at 17 kDa with a 1.9 nm Stokes radius.

B). A standard curve generated elution profile of the standards plotted against their known Stokes radii. The elution volume for peak 1 was 12.8 ml representing a Stokes radius of 5.1 nm and the elution volume for peak 2 was 14.8 ml representing a Stokes radius of 5.1 nm.
