## Supplemental Figure 3 for "Merlin Tumor Suppressor Function is Regulated by PIP_2_-Mediated Dimerization"

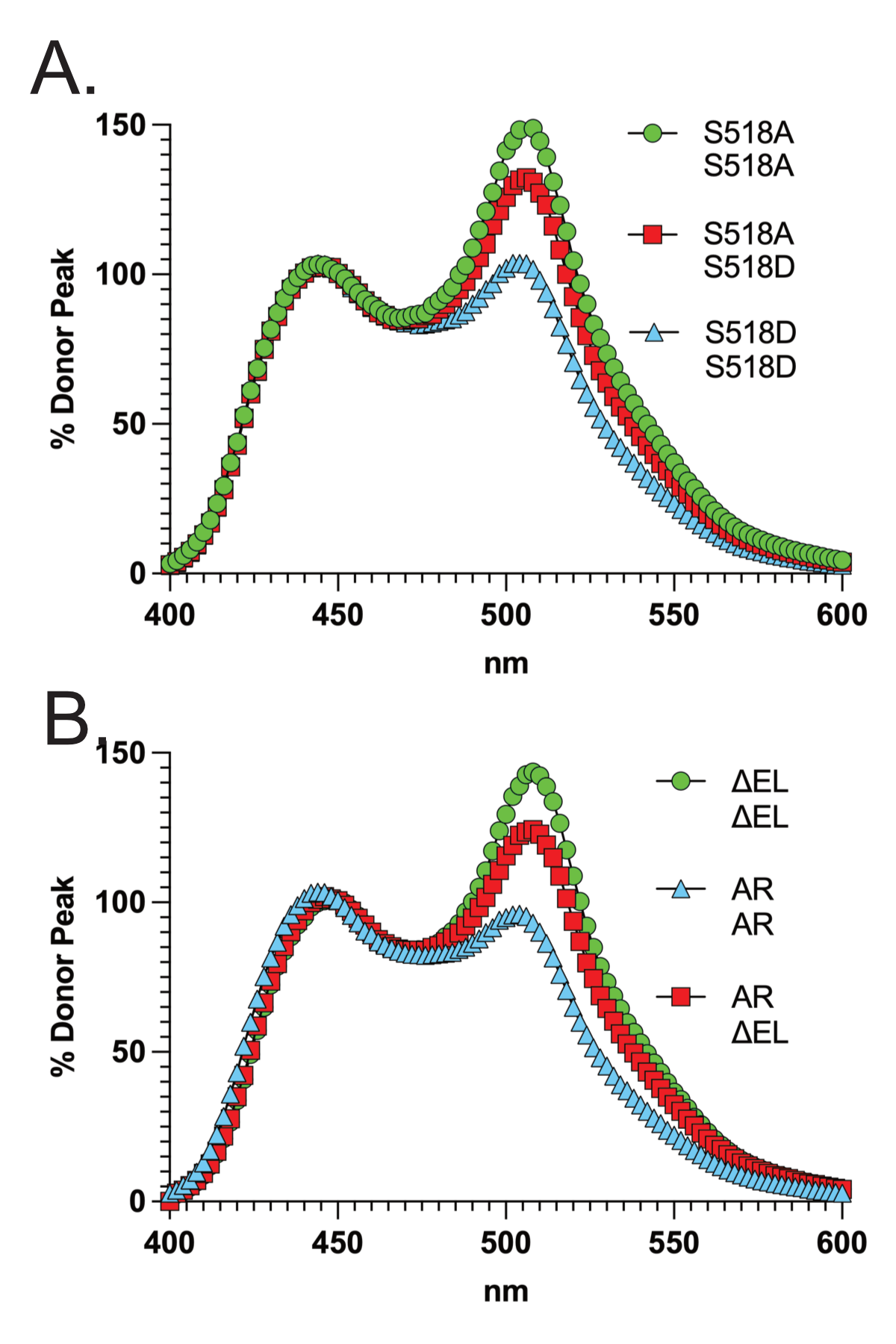

A). BRET assays showing emission spectrum from 400 nm to 600 nm from dimerization assays for combinations of the Merlin phosphorylation mutants S518A:S518A, S518A:S518D and S518D:S518D normalized to the 450 nm peak.

B). BRET assays showing emission spectrum from 400 nm to 600 nm from dimerization assays for combinations of the Merlin conformation mutants AR:AR, AR:ΔEL and ΔEL:ΔEL normalized to the 450 nm peak.
